# Forecasting of phenotypic and genetic outcomes of experimental evolution in *Pseudomonas syringae* and *Pseudomonas savastanoi*

**DOI:** 10.1101/2024.02.10.579745

**Authors:** Jennifer T. Pentz, Aparna Biswas, Bassel Alsaed, Peter A. Lind

## Abstract

Microbial experimental evolution is commonly highly repeatable under identical conditions, indicating a potential for short-term evolutionary forecasting. However, it is unclear to what extent evolutionary predictions can be extrapolated to related species adapting in similar environments, which would enable direct testing of general forecasting models and biological assumptions. To further develop a model system for evolutionary forecasting based on adaptation to static culture conditions, we experimentally tested previous predictions for *Pseudomonas syringae* and *Pseudomonas savastanoi*. In addition to sequence divergence, these species also differ in their repertoire of diguanylate cyclases that can be mutationally activated to produce the adaptive wrinkly spreader (WS) phenotype and genes for biosynthesis of exopolysaccharides. After experimental evolution, we isolated 32 independent WS mutants for *P. syringae* and 37 WS mutants for *P. savastanoi* that had increased ability to colonize the air-liquid interface and reduced motility. As predicted, most mutants had mutations in the *wsp* operon followed by rarer promoter mutations upstream of uncharacterized diguanylate cyclases. Surprisingly, no mutations were found in *wspF,* the most commonly mutated gene in the previously characterized species, which was explained by differences in relative fitness. While prediction of mutated regions was largely successful for WspA, mutations in WspE had a divergent pattern for both species. Surprisingly, deletion of known exopolysaccharide loci previously shown to contribute to the adaptive WS phenotype in other species did not reduce fitness, suggesting the presence of additional adhesive components under c-di-GMP control. This study shows that evolutionary forecasts can be extended to related species, but that differences in the genotype-phenotype-fitness map and mutational biases limit predictability on a detailed molecular level.

**Author summary:** Biological evolution is often observed to be repeatable in the short-term, which suggests that it might be possible to forecast and ultimately steer evolution. Evolutionary processes are fundamental to biology but are also central to major societal problems, including antibiotic resistance, cancer, and adaptation to climate change. Experimental evolution with microbes makes it possible to study evolutionary processes in real-time over many generations to allow direct tests of evolutionary forecasts. However, a fundamental problem is that predictive models are usually based on previous experimental data which limits the novelty of the prediction beyond simple repeatability. A more challenging issue is to predict to what degree similar species evolve in similar ways in similar environments. Here we show that one of the best characterized bacterial experimental evolution model systems, biofilm formation at the surface of static tubes in *Pseudomonas*, can be extended to related species evolving in similar environments. This allowed us to directly test previous evolutionary forecasts to show that similar phenotypes evolved in similar environments, but that predictions of molecular details often fail. This study also elucidates the causes for failed forecasts to allow continuous improvements in predictive models and to delineate the limits of evolutionary forecasting.

## Introduction

The predictability of evolution is a fundamental biological question and the ability to forecast evolutionary events could lead to applications in infectious diseases, cancer, and aid in predicting the ability of organisms to adapt to climate change [1–3]. Traditionally many biologists have been skeptical of the possibility to predict future evolutionary events given that there are often millions of possible mutations and the dependence of previous events, historical contingencies, is difficult to take into account. Adaptive laboratory evolution replay experiments, where multiple replicates of microbial or viral populations evolve under identical conditions, show that evolution is often surprisingly repeatable, suggesting that it might be possible to predict short-term evolutionary outcomes. However, the results from “historical difference experiments”, where replicate lines first evolve separately to evolve different life histories, and then evolve in the same environment shows that the effects of historical contingencies can be significant even in cases where the differences are relatively minor (reviewed in [4]). Although there are many experimental evolution studies exploring the predictability of evolution in terms of repeatability, it is rarely attempted to forecast experimental evolution in the sense of using “data from the past or present to make a prediction about the future” [5] under significantly different experimental conditions [6].

A related question is whether the evolution of related species can be predicted based on previous data and models from another species. To put it simply: will similar species evolve in similar ways in similar environments? This can be seen as an extreme case of an historical difference experiment in that the historical differences have accumulated over millions of years, including gain and loss of genes through horizontal gene transfer. The success of these kinds of evolutionary forecasts will ultimately depend on the conservation of the genotype-to-phenotype map (molecular functions and regulatory interactions) between species, the relative fitness effects of mutations and mutational biases, including mutational hot spots.

We have previously shown that a mathematical model [7] of the genotype-to-phenotype map constructed for *Pseudomonas fluorescens* SBW25 (hereafter *Pflu*) together with experimental data could be used to predict evolutionary outcomes on various biological levels for the related species *Pseudomonas protegens* Pf-5 (hereafter *Ppro*) [8]. This demonstrated that the well-established “wrinkly spreader” (WS) model system could be extended to a related species. The experimental setup of the WS system is simple: Bacteria are grown under static conditions where strong selection for access to oxygen at the surface leads to increased frequency of mutants with the ability to colonize the oxygen-replete air-liquid interface through adhesion between cells and/or the wall of the growth vessel [9]. Wrinkly spreader mutants can be identified by their divergent colony morphology on agar plates and can be easily isolated even when present at a frequency lower than 1%, with most mutants requiring only a single mutation [8,10,11]. WS mutants all have mutations activating production of c-di-GMP by diguanylate cyclases (DGCs), a conserved signal for biofilm formation resulting in increased production of exopolysaccharides that are the main structural components of the biofilm [6,10,11]. While there are many different DGCs that can be activated by mutation to produce the WS phenotype [11], three mutational pathways dominate in both *Pflu* and *Ppro* (Wsp, Aws, and Mws) [7,8,10]. These pathways have a larger mutational target size leading to a hierarchy of genetic routes to WS, where the most commonly used pathways are those under negative regulation followed by more rare promoter mutations [11].

Here we explore if these predictions can be extended to more divergent species with different complements of DGCs and exopolysaccharides. Are more closely related species more likely to evolve in the same way or are evolutionary outcomes more idiosyncratic and dependent on largely unpredictable differences in fitness effects of mutations and mutational hotspots? We test our previously published predictions [8], referred to as P1-P8 below, for two more distantly related *Pseudomonas* species: *Pseudomonas syringae* pv. tomato DC3000 (hereafter *Psyr*) and *Pseudomonas savastanoi* pv. phaseolicola 1448A (hereafter *Psav*):

P1. Mutants with increased ability to colonize the air-liquid interface by increased cell-cell or cell-wall adhesion will evolve and rise to high frequencies.

P2. These mutants will primarily use exopolysaccharides for increased adhesion as they provide the highest structural stability and fitness with cellulose predicted to be used for *Psyr* and Psl for *Psav*.

P3. Mutants will have reduced motility due to increased production of c-di-GMP by DGCs, which is a signal for increased production of exopolysaccharides and give the mutants a WS morphotype.

P4. Most mutations will cause loss of function, followed by promoter mutations and even less frequent activating mutations and double inactivating mutations.

P5. Mutations in the molecular networks of the negatively regulated DGCs WspR (for *Psyr* and *Psav*), followed by MwsR (for *Psyr* only) will be the most common route to WS due to a large mutational target size.

P6. A previously developed mathematical model [7,12] predicts that most mutations will be in *wspA*, *wspE*, *wspF* for *Psyr* and *Psav*, and *mwsR* for *Psyr* only.

P7. Mutated regions for proteins in the Wsp and Mws pathways can be predicted based on previous WS mutations in *Pflu* and *Ppro*.

P8. Mutations will not be evenly spread between the predicted *wspA*, *wspE*, *wspF* and, *mwsR* genes with large mutational targets, but found in those where mutations have the largest beneficial fitness effects.

Experimental tests of our predictions confirmed that WS mutants with increased ability to colonize the air-liquid interface and reduced motility commonly evolved for both *Psyr* and *Psav*. Most WS mutations were in regions predicted to disrupt negative regulation of DGCs followed by more rare promoter mutations for both species. As predicted, Wsp was the most commonly mutated pathway for both species followed by Mws in *Psyr*. However, the mutational patterns in *Psyr* and *Psav* differed from those found in *Pflu* and *Ppro,* which could partly be explained by fitness differences. Surprisingly, deletion of known exopolysaccharide biosynthesis genes in WS mutants did not reduce fitness for either species, suggesting that alternative or additional adhesion factors are used. Mutational hot spots apparent in *Pflu* and *Ppro* were not conserved, but the most common mutation was identical for *Psyr* and *Psav*. This study shows that it is possible to make evolutionary forecasts over larger phylogenetic distances, but that unpredictable differences in mutational hotspots and fitness effects of mutations limits detailed genetic predictions.

## Results

### WS mutants were isolated after experimental evolution with *Psyr* and *Psav*

To examine the causes and limitations of evolutionary predictability across longer phylogenetic distances, we chose *Psyr* and *Psav* that are further diverged from *Pflu* than *Ppro*, with 75% average amino acid identity in genes commonly targeted by WS mutations compared to 81% in *Ppro* [8]. While *Psyr* and *Psav* are more closely related to each other than *Pflu* and *Ppro*, there is still substantial sequence divergence between them [*wspA* (89% nucleotide identity, 96% amino acid identity), *wspE* (85% nt, 90% aa) and *wspF* (84% nt, 93% aa)].

We performed experimental evolution in static growth microcosms for 7 days at 25°C to select for wrinkly spreader (WS) mutants that colonize the air-liquid (AL) interface in both *Psyr* and *Psav*. As predicted (P1), colonization of the AL interface was observed, and 32 independent WS mutants were isolated for *Psyr* and 37 mutants for *Psav* based on their divergent colony morphology on agar plates.

### WS phenotypes are caused by mutational activation of DGCs with Wsp being the most commonly used pathway

To further evaluate our predictions, we first determined the genetic cause of WS evolution for all isolated mutants using a combination of Sanger and Illumina sequencing. This allowed us to find mechanistic clues for increased adhesion, including if c-di-GMP production is involved (P3), as well as to evaluate our genetic predictions in terms of types of mutations (P4), DGC pathways (P5), mutated genes (P6) and mutated regions (P7). Compared to *Pflu*, *Psyr* and *Psav* exhibit adifferent repertoire of available DGCs and EPSs [8]. Importantly, they do not encode all three common genetic pathways (Wsp, Aws, MwsR) to WS used in *Ppro* [8] and *Pflu* [10] where they account for approximately 99% of WS mutants [7]. *Psyr* encodes Wsp and MwsR (but not Aws), while *Psav* only encodes Wsp. Despite these major differences, most mutations were found in the common pathways available, with 30/32 mutations occurring in Wsp and Mws (Figure 1A) for *Psyr*, and 34/37 mutations occurring in Wsp (Figure 1B) for *Psav* (P5).

**Fig 1.**
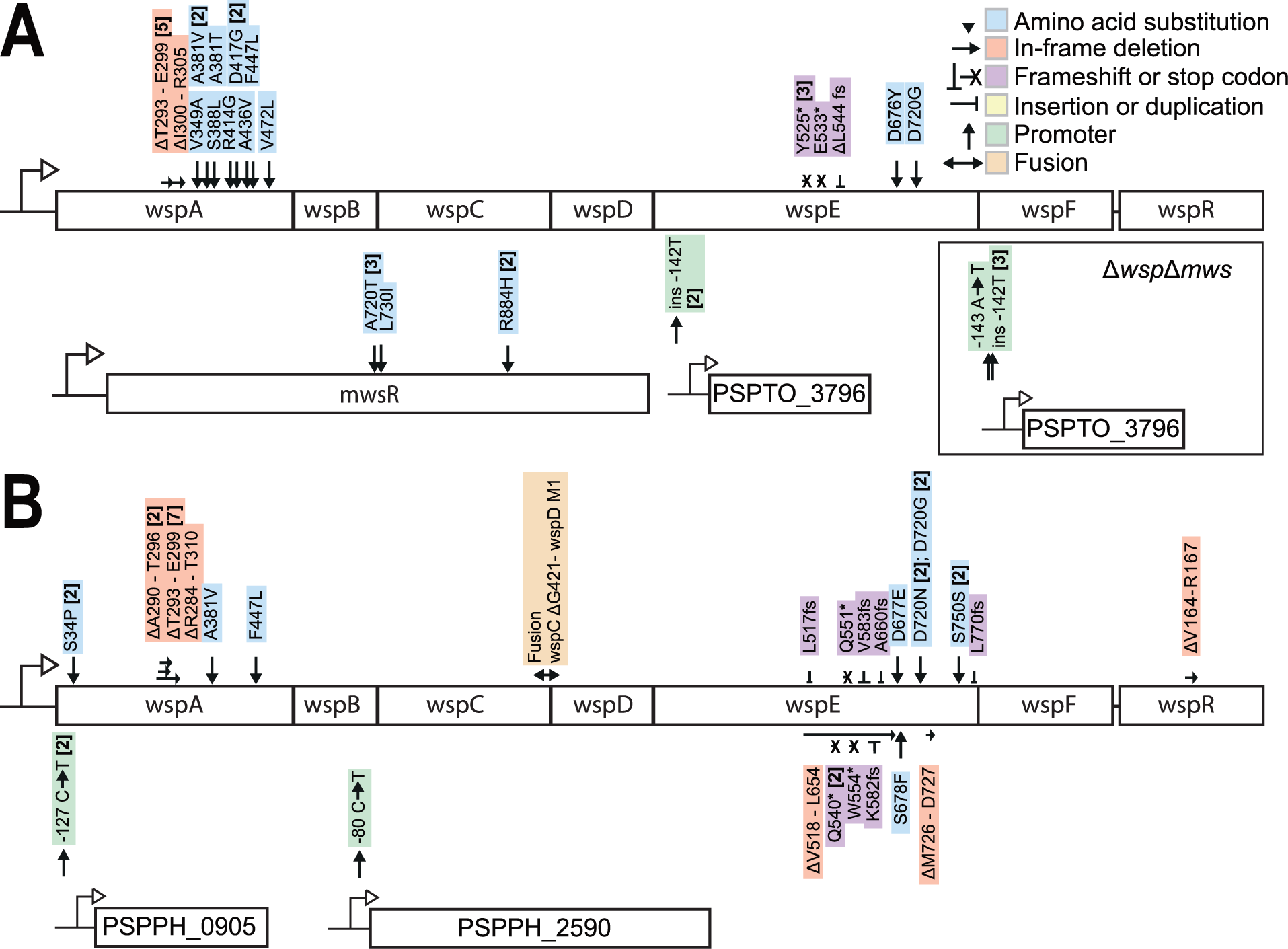
Mutations identified in WS mutants. (*A*). Thirty-two independent WS mutants were isolated for *Psyr*. Four additional WS mutants were identified after experimental evolution with Δ*wsp* Δ*mws* double deletion mutant. (*B*). Thirty-seven independent mutants were isolated for *Psav* after experimental evolution. Numbers in brackets are the number of independent mutants found. Details are available in S1 Table.

As predicted most mutations were expected to disrupt negative regulation of DGCs followed by rarer promoter mutations (P4). In *Psyr*, two mutations were found upstream of PSPTO_3796, encoding a putative DGC predicted to be localized to the cytoplasmic membrane. To determine if there are also other DGCs that can be activated by mutation, as previously demonstrated in *Pflu* [11], *wspABCDEFR* and *mwsR* were deleted in *Psyr*, followed by experimental evolution using an identical protocol as for the ancestor. This allowed isolation of four independent WS mutants that all had mutations in the PSPTO_3796 promoter indicating a lack of alternative pathways or a high mutation rate at this promoter. Similarly, in *Psav*, two mutations occurred upstream of PSPPH_0905, encoding a putative DGC, and one mutation was found upstream of PSPPH_2590, encoding a putative protein with both a DGC domain and a putative phosphodiesterase domain. Both PSPPH_0905 and PSPPH_2590 were predicted [13] to be localized to the cytoplasmic membrane. Thus, all WS mutants for both *Psyr* and *Psav* have mutations linked to DGCs presumably increasing production of c-di-GMP (P3).

### Divergent mutational patterns in Wsp

Despite the similarities in mutated pathways and types of mutations, some aspects of the mutational patterns were strikingly different when compared with the previously isolated *Pflu* and *Ppro* WS mutants (S1 Figure). These dissimilarities could be due to differences in the genotype-to-phenotype map (similar mutations do not produce a similar phenotype) or differences in fitness (the relative fitness effects of mutations in specific genes are not conserved) (P7 and P8). The most striking difference was the lack of mutations in *wspF*, which is surprising as it encodes the negative regulator of the Wsp network and any inactivating mutation is expected to result in a WS phenotype and it is the most commonly mutated gene in both *Pflu* and *Ppro* WS mutants [8,10]. Another marked difference is that there are many mutations in *wspA* (16 for *Psyr* and 14 for *Psav*), which has a high predicted mutation rate to WS [7], but where mutations were not found after experimental evolution for *Pflu* and *Ppro* [8,10]. This might be explained by an apparent shared mutational hot spot in *wspA* accounting for 16% of mutations for both *Psyr* and *Psav* that results in a T293 - E299 in-frame deletion between direct repeats of 7bp in *Psav* and 9 bp in *Psyr*. This mutation is predicted to lead to loss-of-function of the methylation sites in the region 280-310 [8] thereby disabling the functional interaction with negative regulator WspF. Additionally, this pattern could be explained by differences in relative fitness as WspA mutants in *Pflu* and *Ppro* have low fitness and are outcompeted by other WS mutants [7,8].

A third clear difference in mutational patterns is the types of mutations observed in *wspE*, which is also predicted to be a commonly mutated gene [7]. Mutations in *wspE* have previously been found in both *Pflu* [7,10] and *Ppro* [8], but they are in all cases missense mutations near the phosphorylation site (4-aspartyl phosphate at D720) in the response regulatory domain. These mutations were predicted [8] to lead to loss-of-function of the phosphorylation reaction that activates WspF leading to disabling of negative regulation and constitutive activation of WspR. For both *Psyr* and *Psav*, missense mutations in the same region are also found, but there were many different insertion and deletion mutations, including frameshifts and nonsense mutations causing loss of the entire CheW-like and response regulatory domains (S2 Fig). Thus, mutated regions could not be predicted based on previous data from *Pflu* and *Ppro* for the WspE protein (P7, S2 Fig).

### Reconstructed WS mutants have reduced motility and colonize the air-liquid interface

We reconstructed a subset of mutants representing the diversity of mutations for *Psyr* and *Psav*, creating two independent identical mutants in ancestral strains expressing either of two fluorescent proteins (YFP and BFP). This ensures that the identified mutation is the sole cause of the phenotypic changes observed and that there are no secondary mutations influencing the results. We also constructed strains with mutations in *wspF* and PSPTO_0339 (*dgcH*) that were predicted to be observed at high frequency [8] based on previous data [11] but were not found after experimental evolution. This set of mutants were used for phenotypic characterization and fitness measurements with the aim to further examine the causes of the observed mutational patterns.

The colony morphology for the reconstructed WS mutants were most pronounced for *wspA* and *wspE* mutants, but the wrinkly phenotype was less characteristic than for WS mutants in *Pflu* and *Ppro* (Fig 2A, Fig 2C). Clear colonization or air-liquid interface was observed for most mutants (P1), but the *wspF* and PSPPH_0905 mutants did not form thick biofilms (Fig 2A, Fig 2C). Swimming motility was clearly reduced for most mutants (Fig 2B, Fig 2D) (P3), but not for *wspF* for both *Psyr*/*Psav* and PSPPH_0905, suggesting a connection between ability to colonize the air-liquid interface and reduced motility. For *Pflu* and *Ppro*, *wspF* mutants are the most commonly isolated after experimental evolution and they have reduced motility and a strong wrinkly colony morphology. This suggests that the WS genotype-to-phenotype map is not conserved between *Pflu*/*Ppro* and *Psyr*/*Psav*.

**Fig 2.**
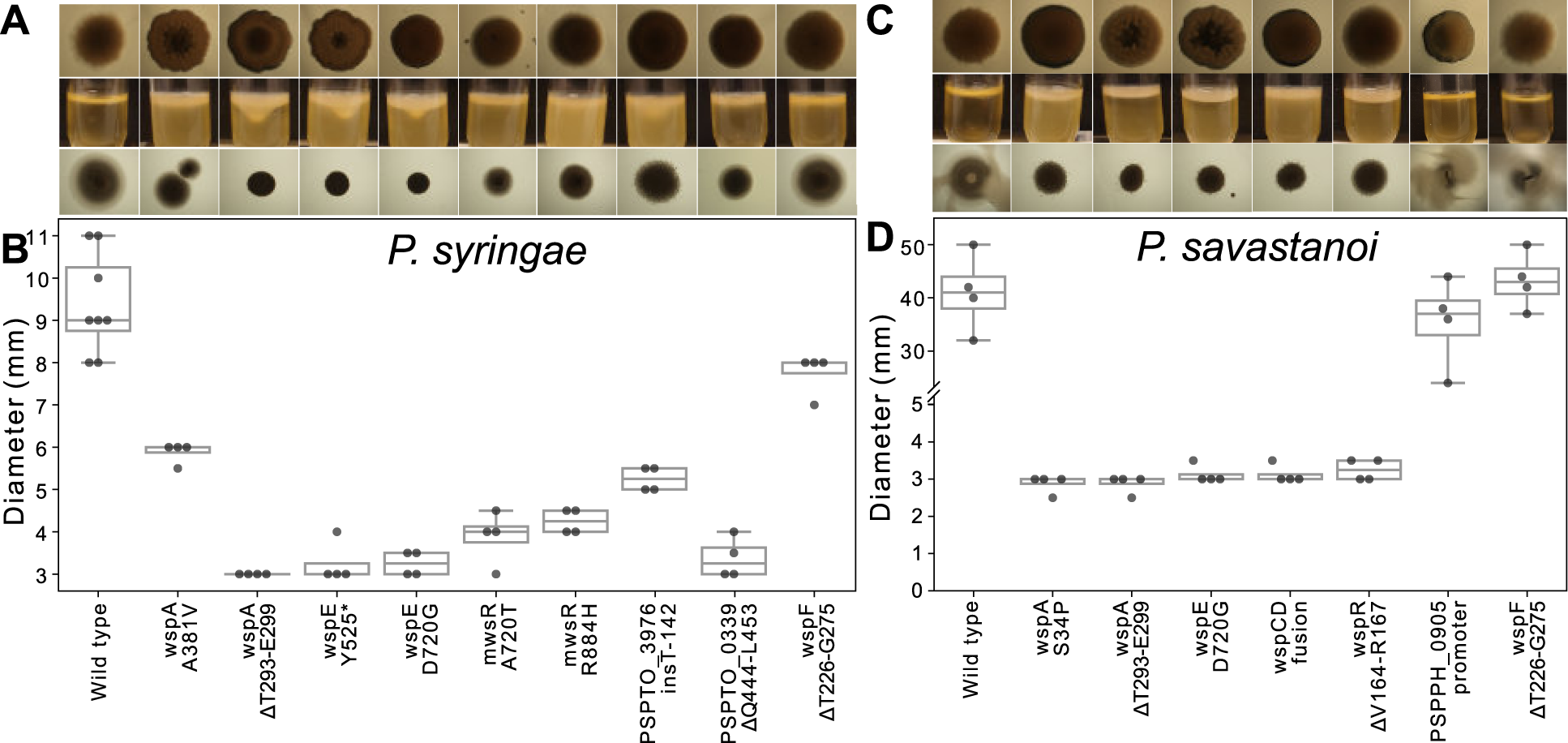
Phenotypic characterization of reconstructed mutants. (*A*). *Psyr* colony morphology (1.5% agar), biofilm formation under static growth, and swimming motility (0.3% agar) of reconstructed mutants. (*B*) Swimming motility of *Psyr* reconstructed mutants. (*C*) *Psav* colony morphology (1.5% agar), biofilm formation under static growth, and motility (0.3% agar) of reconstructed mutants. (*D*) Swimming motility of *Psav* reconstructed mutants.

### Most WS mutants have similar competitive fitness except for less fit WspF mutants

To determine how differences in fitness between WS mutants could explain the observed mutational patterns, we conducted two types of competitive fitness assays, similarly to previous studies [7,8,11,14]. The first assay, measuring what we refer to as “invasion fitness”, measures competition of a WS mutant starting at 1% against the wild-type ancestor at 99% and is intended to measure fitness at the early stages of air-liquid interface colonization. In the other assay, measuring “competition fitness”, each WS mutant is competed 1:1 against a *wspA* reference WS mutant that represents the most common mutant isolated after experimental evolution for both *Psyr* and *Psav*. These assays can provide data to explain the absence of expected mutants, like WspF (P6), after competition and determine if rare promoter mutants have similar fitness to the commonly mutated pathways so that mutational target size is a main determinant of evolutionary outcomes (P4) [11].

All WS mutants had similar invasion fitness against the ancestor and the reference strain (WspA ΔT293-E299) except for *Psyr* WspF that is substantially lower, but still significantly positive (*p*=0.002; one-way ANOVA, *F_9,86_*=16.7, *p*<0.0001, pairwise differences assessed with Tukey HSD) (Fig 3A, Fig 3C). Thus, all WS mutants are expected to be able to rapidly increase in frequency when appearing in an ancestral population.

**Fig 3.**
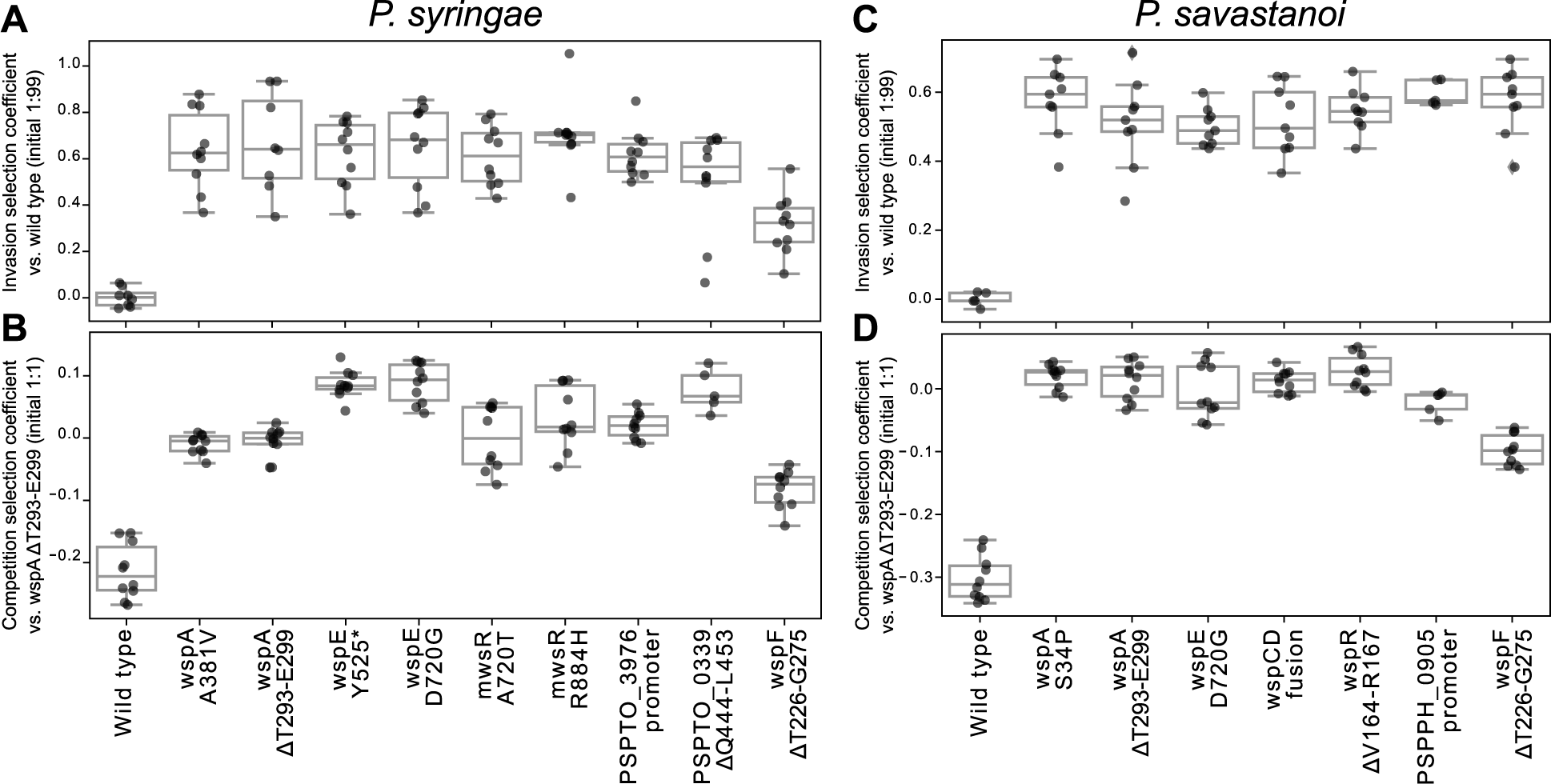
Fitness of reconstructed *Psyr* and *Psav* WS mutants. Invasion fitness (A, C) of mutants was measured against the wild-type (initial ratio 1:99 mutant:wild-type) while competition fitness (B, D) was measured against the most common WspA mutant (initial ratio 50:50). Reciprocal pairwise competitions were performed in YFP and BFP backgrounds.

The competition assay showed that in *Psyr* both WspE mutants had higher fitness than the WspA reference (Fig 3B; WspE Y525* *p*<0.0001, WspE D720G *p*<0.0001; one-way ANOVA, F_9,85_=68.5, *p*<0.0001, pairwise difference assessed with Tukey HSD). Surprisingly, the PSPTO_0339 mutant also had higher fitness (Fig 3B; *p*<0.0001) despite it not being observed after experimental evolution, which could possibly be explained by a smaller mutational target size leading to a lower mutation rate to WS [11]. The other *Psyr* WS mutants had similar fitness as the WspA reference with the exception of the WspF mutant, explaining why WspF mutants were not observed after experimental evolution (Fig 3B). The WspF mutant also had clearly reduced competition fitness for *Psav* (Fig 3D) while the other WS mutants had similar fitness as the WspA reference, with the PSPPH_0905 mutant showing a small but insignificant decrease (WspF *p*<0.0001, one-way ANOVA, F_7,67_=149.1, pairwise differences assessed with Tukey HSD). These results strongly suggest that the lack of WspF mutants after experimental evolution is due to their lower fitness and that mutations are found clustered in the genes where mutations have the largest beneficial fitness effects (P8).

### Deleting exopolysaccharide biosynthesis genes in WS mutants do not reduce fitness

To test the prediction that mutants with increased ability to colonize the air-liquid interface would use exopolysaccharides as a key adhesive structural component (P2), we used Tn5 transposon mutagenesis of WS mutants to isolate insertion mutants that reverted to the ancestral colony morphology. In *Pflu*, WS mutants primarily rely on increased production of cellulose [14,15]. *Psyr* also encodes the genes for cellulose biosynthesis, so we predicted that Tn5 insertions would mainly be found in the *wss* operon [8]. After Tn5 mutagenesis of WspA and WspE mutants of *Psyr* we determined insertion sites and we found insertions in *wssA*, *wssB*, *wssC*, *wssE*, *wssF* and *wssI* as predicted, strongly suggesting that the divergent colony morphology is caused by increased production of cellulose.

As *Psav* does not encode genes for biosynthesis of cellulose, Pga, or Pel that have been shown to be important for WS mutant fitness in SBW25 and Pf-5 [8,14], Psl was predicted [8] to be used as the main structural component based on its role in *Pseudomonas aeruginosa* [16] and *Pseudomonas simiae* [17]. We did a similar Tn5 mutagenesis screen for *Psav* WspA mutants and found insertions in the alginate operon genes *algI*, *algX*, *alg44*, *alg8* that reverted the WS mutants back to the ancestral colony morphology, which instead suggest that increased alginate production is the main cause of the divergent colony morphology.

While the Tn5 mutagenesis experiments indicates that increased production of cellulose in *Psyr* and alginate for *Psav* is responsible for the wrinkly colony morphology in Wsp mutants, it is not necessarily the main structural component required for successful air-liquid interface colonization leading to increased fitness. Therefore, we deleted the *wss* operon from several WS mutants of *Psyr* and the *alg* operon from WS mutants of *Psav* to determine the roles of these exopolysaccharides in biofilm formation under static conditions. Surprisingly neither of these deletions reduced fitness in the invasion assay or the competition assay (Fig 4A). We also constructed strains of WS mutants in *Psav* with the *psl* operon deleted or both *alg* and *psl* operons deleted, but no reduction in either invasion or competition fitness was observed (Fig 4B). These results suggest that these exopolysaccharides, although apparently increasingly produced in WS mutants, do not contribute significantly to increased fitness or that in their absence other unknown adhesive factors can substitute for their function.

**Fig 4.**
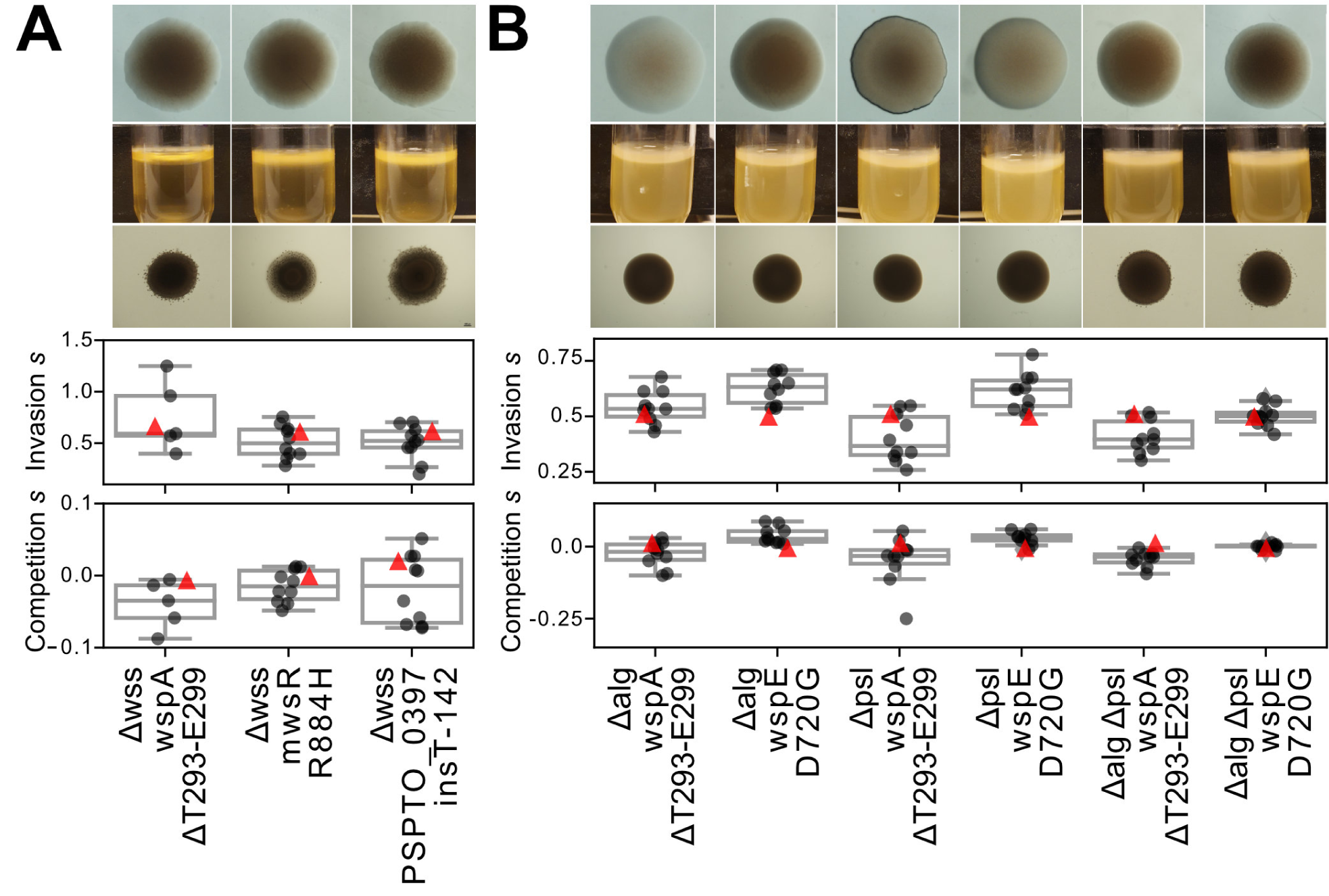
Deletion of known exopolysaccharides biosynthesis genes does not decrease fitness. (*A*) *Psyr* WS mutants with deletion of the *wss* operon. From top: colony morphology (1.5% agar), biofilm formation under static growth, and swimming motility (0.3% agar) of reconstructed mutants. Invasion and competition fitness of identical WS mutants with the *wss* operon intact shown as red triangles for comparison (from Fig 3). (*B*) *Psav* WS mutants with deletion of the *alg* and/or *psl* operons. Colony morphology (1.5% agar), biofilm formation under static growth, and swimming motility (0.3% agar) of reconstructed mutants. Invasion and competition fitness of identical WS mutants with the *alg* and *psl* operons intact shown as red triangles for comparison (from Fig 3).

To further investigate the molecular basis of biofilm formation in WS mutants devoid of cellulose production for *Psyr* and psl and alginate production for *Psav* we used proteinase K that cleaves peptide bonds, cellulase that cleaves β-1-4 glycosidic bonds present in some polysaccharides or DNAse that cleaves phosphodiester linkages in DNA. After 48 h of incubation with proteinase K a reduction in biofilm formation at the air-liquid interface was apparent for cultures of *Psyr* Δ*wss* WspA ΔT293-E299 and to a lesser degree for *Psyr* Δ*wss* MwsR R884H, which could indicate that an adhesive protein or peptide plays a major structural role (S3 Fig). However, addition of proteinase K only delayed biofilm formation and at 72 h it was clearly visible at the air-liquid interface for both mutants (S3 Fig). None of the three enzymes reduced biofilm formation in *Psav* Δ*alg* Δ*psl* WspA ΔT293-E299 (S3 Fig). Addition of cellulase to *Psyr* and *Psav* cultures increased growth both at the air-liquid interface and in the broth phase, which is presumably an artifact caused by degradation of sugars and oligosaccharides in the growth medium increasing the concentration of monosaccharides to speed up growth.

## Discussion

Explicit testing of short-term evolutionary forecasts allows an unbiased view into the factors that govern evolutionary processes and critical examination of fundamental biological assumptions. Here we show that short-term evolutionary forecasts are to some degree extendable to similar species evolving in similar environments, but that differences between species can make evolutionary outcomes idiosyncratic and unpredictable on the molecular level. Several of our previously published predictions [8] were successful, including the evolution of mutants with increased ability to colonize the air-liquid interface (P1) and the mutational activation of c-di-GMP production resulting in reduced motility (P3). Interestingly, the mutants with higher motility (Fig. 2) have lower fitness (Fig. 3) in competitions strengthening a link between reduced motility and fitness. The mutational pattern of common loss-of-function mutations, resulting in loss of a key function of the protein, and more rare promoter mutations (P4) was observed in all four species tested so far [8,11], suggesting that the mutational target size is a major determinant of evolutionary outcomes in this model system where multiple genetic pathways to similar high-fitness phenotypes are available. This is also reflected in the conservation of the order of frequency of pathways used (P5) for all four species tested (this work and [8,10]) that has also been found in *Pseudomonas simiae* PICF7 [17] that is more closely related to *Pflu*. Yet, on a more detailed genetic level there is less conservation between the different species, but with indications that more closely phylogenetically related species evolve more similarly even when they encode different exopolysaccharide structural components.

At the gene level there is less overlap with the previous studies in *Pflu*/*Ppro*, where mutations in the *wsp* operon were found primarily in *wspF*, while for *Psyr*/*Psav* mutations no mutations were found in *wspF* and most mutations were in *wspA* and *wspE*. This aligns well with P6 that mutations will be unevenly spread between the genes predicted by a previously developed model of the molecular networks [7] and clustered in the genes where mutations can provide the largest beneficial fitness effects. We confirmed that fitness effects provide a reasonable explanation for these patterns in all four species ([7,8] and this work) and it might also explain observations in the even more distantly related *Burkholderia cenocepacia* where only mutations in *wspA* and *wspE* were found [18]. The weak colony morphology phenotype, high motility and low fitness of the WspF loss-of-function mutants in *Psyr* and *Psav*, suggests that the Wsp regulatory network in these species are clearly different from that in *Pflu* and *Ppro*. In contrast, partial loss-of-function mutations in interacting proteins WspA and WspE were found and produced strong WS types, as predicted. One possible explanation for this is that there is an unknown protein partially compensating for the absence of WspF methylesterase activity in *Psyr* and *Psav*, something that would be very difficult to predict *a priori*. Another example of differences in fitness effects are WS mutations in WspR that are expected to be relatively frequent [7,19] but that are not found after experimental evolution in *Pflu* as they have much lower fitness [7], while in *Psav* the WspR mutant had similar fitness as other WS mutants (Fig 3C, Fig 3D). Clearly an inability to predict which genes will harbor mutations that leads to the most fit mutants will limit the success of evolutionary forecasts, but experimental data for a small number of constructed mutants might suffice to identify those genes without knowledge of the full distribution of fitness effects for mutations for each gene.

Relative fitness of WS mutants is not necessarily the only reason for the differences in mutational patterns between species. The difference in the mutation spectrum in WspE hints that its molecular function or interactions are in some way different between *Pflu*/*Ppro* that only had very specific point mutations around an active site, and *Psyr*/*Psav* that also had mutations causing loss of the entire CheW-like and response regulatory domain by a range of different indels and point mutations. This presumably means that the number of possible WS mutations in *Psyr* and *Psav* is higher, which could bias the observed mutational patterns towards *wspE*. Thus, the mutated regions and types of mutations for WspE could not be predicted based on previously observed mutations in other species (P7, S2 Fig). In contrast, there was a strong similarity between mutations in WspA in *Pflu* [7] and in *Psyr*/*Psav* even if exact molecular parallelism was rare (S1 Fig).

The most obvious forecasting failure is the limited role played by the exopolysaccharides cellulose for *Psyr* and alginate and psl for *Psav* (P2). Although it seems likely that the exopolysaccharides are indeed produced at a higher level in WS mutants, based on the difference in colony morphology of deletion mutants, there was no reduction in fitness either in invasion or competition assays. In *Pflu* the deletion of the cellulose operon in WS mutants causes a major reduction in competitive fitness although the alternative Pga exopolysaccharide can partially compensate to allow air-liquid colonization [14]. The Pga biosynthesis genes are not present in either *Psyr* or *Psav*, but it is likely that there are unidentified adhesive factors, possibly including uncharacterized exopolysaccharides [8], that are also under regulation of c-di-GMP in these species. The ability of proteinase K to delay biofilm formation in *Psyr* WS mutants with the cellulose biosynthesis operon deleted suggests that a protein or peptide plays a role in increasing the ability to colonize the air-liquid interface. However, after extended incubation there is biofilm formation exceeding that of the ancestor indicating that adhesive factors under c-di-GMP control resistant to degradation by proteinase K is still produced. In *Psav*, neither cellulase, proteinase K or DNase I visibly reduced biofilm formation in the WspA mutant with the *psl* and *alg* operons deleted, which could point to complex adhesion mechanisms that is not disrupted cleavage of a single type of chemical bond.

By repeatedly conducting experimental tests of forecasts of WS evolution for different species there are opportunities to use new data to iteratively improve forecasts. The mathematical model used in Lind *et al.* [7] that formed the basis for the predictions [8] on the gene and pathway levels only included the regulatory connections of the main DGC networks with a simplified assumption that disabling mutations, reducing reaction rates, were ten times more common than enabling mutations. It might be possible to improve this model with more experimentally informed probabilities of mutations affecting mutation rates and change the models of regulatory networks considering an increased understanding of the molecular functions of the proteins involved. An alternative model that might be more suitable for forecasting simple one-step evolution from a smooth ancestor to a WS phenotype is to use the mutational patterns accumulated by experiments in different species (S1 Fig) to estimate the mutational target size for each gene, in terms of the number of possible WS mutations, without any consideration of the mechanistic details of the regulatory networks. The success of such a model is expected to be limited by the presence of mutational hot spots that have been observed to some degree in all species tested so far. In *Pflu*, just two mutations account for 42% of the total mutation rate to WS [7], a single mutation accounts for 26% of WS mutants observed in *Ppro* and the same mutation in *wspA* accounts for 16% of mutants in both *Psyr* and *Psav*. Thus, even though there are likely hundreds of possible WS mutations (S1 Fig, [7,8,10,11,20]), just a few account for a major part of the genetic variation produced in the populations increasing the probability of parallel molecular evolution for each species.

Clearly, differences in mutation hot spots can bias evolutionary outcomes but it is not obvious to what extent this will limit predictability and how much variation we should expect in evolutionary outcomes for different species with a conserved genotype-to-phenotype map. Sun and Lind [21] addressed this problem by combining a target size model of the WS model system with a distribution of mutation rates, that the mutation rate of each possible WS mutation is sampled from, to examine how much variation in mutation rates would affect evolutionary outcomes on the levels of individual mutations, genes, and pathways. Based on repeated observations that mutational hot spots can have a major effect on evolutionary outcomes [7,8,14,22–26], it is clear that a distribution of mutation rates must have a long-tail of a small fraction of mutations that have greatly increased mutation rates and a log-normal distribution was used to examine how mutational biases can limit evolutionary predictability [21]. The theoretical predictions from Sun and Lind aligns well with the experimental data for the different *Pseudomonas* species ([7,8,10] in that it predicts that even with inclusion of strong mutational hot spots, evolution on the pathway level is predictable between species (P5) so that Wsp is typically the most commonly mutated pathway.

Another aspect that can also be explained by the distribution of mutation rates model is why the rare pathways used for the different species are likely to be different. Most DGCs are not under negative regulation and they must be activated by mutations in promoter regions or specific intragenic activating mutations to produce WS mutants [11]. Given that there are many putative DGCs for each species (39 in *Pflu*, 39 in *Ppro*, 35 in *Psyr* and 34 in *Psav*), which rare DGCs are mutated for each species will determined by the mutation rate of a few nucleotides in promoter regions or in the coding regions, which when sampling from a distribution of mutation rates can make evolution idiosyncratic for pathways with small mutational target sizes [21]. Even though at least nine different DGCs in *Pflu* can be activated by promoter mutations or specific intragenic mutations [11] and seven of these are present in both *Psyr* and *Psav* [8] they were not found to be mutated here. Instead for *Psyr* the promoter of the DGC encoded by PSPTO_3796, that is not present in the other species, was repeatedly mutated. For *Psav,* promoter mutations were found upstream of the putative DGCs encoded by PSPPH_0905 and PSPPH_2590 that both have orthologues in *Psyr*, but not in *Pflu* or *Ppro.* Thus, although the general prediction that promoter mutations will be the second most common type of mutations (P4) is often successful, it is not possible to predict which DGCs those promoters will belong to, as this is likely determined by the mutation rates of a few nucleotides in the promoter region of each DGC. Other limitations of using target size models include the scarcity of suitable experimental data to estimate the distribution of mutation rates and that the genotype-to-phenotype map may not be conserved resulting in large differences in mutational target sizes between species, as observed for WspE in *Pflu*/*Ppro vs*. *Psyr*/*Psav*.

Our results show that short-term evolutionary forecasting in simple environments is possible at higher biological levels even for species that are estimated to have diverged millions of years ago [27] and that have gained and lost significant parts of their genomes. We find that at the lower biological levels of individual genes and mutation evolution is less predictable but still highly repeatable within a single strain. By demonstrating that the well-established wrinkly spreader model system can be extended to even more divergent *Pseudomonas* species we see great promise in continuing to develop this model system that could allow direct testing of evolutionary forecasts across the hundreds of known diverse *Pseudomonas* species [28]. While many of the most basic predictions (P1-P4) could turn out to be generally applicable it will be of particular interest to also see the exceptions to the rules, especially if the underlying causes can be elucidated instead of simply referring to undefined historical contingencies. These forecasting failures can provide fundamental insights into the causes underlying the predictability of evolutionary processes, especially when combined with construction of expected mutants not observed after experimental evolution. These insights can then be used to inform and critically evaluate machine-learning models for evolutionary forecasting that can easily incorporate factors like gene content and phylogenetic relationships but where mutational hotspots can introduce biases in training data and a few nucleotide sequence differences can drastically alter evolutionary outcomes [22].

## Materials and methods

### Strains and media

We used *Pseudomonas syringae* pv. tomato DC3000 (*Psyr*, GenBank: genome AE016853.1, plasmids AE016854.1 and AE016855.1 [29] and *Pseudomonas savastanoi* pv. Phaseolicola 1448A (*Psav*, Genbank genome CP000058.1, plasmids CP000059.1 and CP000060.1, [30]) and its derivatives for all experimental evolution and phenotypic characterization. Cloning of PCR fragments was performed using *E. coli* DH5α *λpir*. *Psyr* and *Psav* were cultured in tryptic soy broth (TSB) supplemented with 10 mM MgSO4 and 0.2% glycerol (TSBGM) for experimental evolution and fitness assays. Genetic engineering was performed using Lysogeny broth (LB), while counter-selection of the *sacB* marker was performed using LB without NaCl supplemented with 8% sucrose. Solid media consisted of 1.5% agar added to LB or TSBGM with or without 10 mg/l Congo red. Motility assays were conducted in 0.3% agar TSBGM. We used gentamicin (10 mg/l) and kanamycin (50mg/l) for *E. coli*, *Psyr* and *Psav,* and nitrofurantoin (50 mg/l) was used to inhibit growth of *E. coli* donor and helper cells after conjugation. All strains were stored at −80°C in LB with 10% DMSO.

### Experimental evolution

For experimental evolution, 1 mL of TSBGM was inoculated with about 10^3^ cells from independent overnight cultures and grown in a deep well plate (1.1 ml, round walls, polypropylene, Axygen Corning Life Sciences) at 25°C for seven days, without shaking. To reduce possible edge effects due to increased evaporation, the wells at the plate’s edges were left unused and served as contamination controls. To sample cells, a 1 µl plastic loop was utilized to collect cells from the bottom, edges, and surface of the air-liquid interface, which were then transferred to an Eppendorf tube containing LB, and vortexed vigorously. In total 100 replicate populations of *Psyr* and 90 replicate populations of *Psav* were sampled to collect a minimum of thirty WS mutants for each species. After seven days of incubation, suitable dilutions were plated on TSBGM plates supplemented with Congo red and incubated for another 48 hours at 25°C. To identify colonies with WS morphology, the plates were screened, and one colony per well was randomly selected based on its position on the agar plate. A total of 32 independent WS mutants for *Psyr* and 39 WS mutants for *Psav* were obtained. Note that this sampling method does not necessarily reflect the frequency of mutants for the entire population and lack of observed WS mutants on the agar plate does not mean that they are not present in the population at a lower frequency than about 1%. The WS mutants were streaked for single cells twice before overnight growth in LB and freezing. The same protocol was used for the *Psyr Δwsp Δmws* strain where four additional mutants were isolated from 30 replicate populations.

### Genetic engineering

Seven *Psyr* WS mutations and six *Psav* WS mutations representing all mutated genes were reconstructed in the wild type strains constitutively expressing either a YFP or BFP marker for use in fitness assays and to confirm the role of the mutation in causing the adaptive phenotype. We also introduced a *wspF* ΔT226-G275 mutation previously found to cause a high fitness WS phenotype in *Pseudomonas fluorescens* SBW25 [7] in *Psyr* and *Psav* fluorescent strains and a PSPTO_0339 ΔQ444-L453 mutation previously found in *Ppro* (corresponding to Q493-L503) [8]. This was accomplished by introducing the mutations or deletion constructs into the wild-type *Psyr* or *Psav* using a two-step allelic replacement protocol. The protocol involved the utilization of the mobilizable pEX18Gm suicide plasmid (GenBank AF047518.1) and was carried out as previously described [8] except at 25°C and without heat shock of recipient strains. Deletion of PSPTO_1026-PSPTO_1034 (Δ*wssABCDEFGHI*) in *Psyr* and deletion of PSPPH_1107-1118 (Δ*algABCDEFGHIJKLX*) and PSPPH_3222-3232 (Δ*pslABCDEFGHIJK*) in *Psav* were also done using the same method. DNA fragments for the mutation of *wspF* in *Psyr* and *Psav* and for deletion of mwsR (PSPTO_4631) in *Psyr* were made by gene synthesis (Thermo Fischer Scientific, GeneArt). DNA fragments for deletion of the *wspABCDEFR and wssABCDEFGHI* in *Psyr* and deletion of *algABCDEFGHIJKLX* and *pslABCDEFGHIJK* in *Psav* was prepared using SOE-PCR to generate fragments surrounding the operons as previously described [8,14].

Strains of *Psyr* and *Psav* expressing fluorescent proteins were constructed using a mini-Tn7 transposon that allows site-specific integration downstream of the *glmS* gene. pUC18R6K-mini-Tn7T-Gm, expressing sYFP2 (KM018300.1) or mTagBFP2 (KM018299.1) from a J23101 promoter (https://parts.igem.org/Part:BBa_J23101) and a BBa_B0034 ribosomal binding site (http://parts.igem.org/Part:BBa_B0034) was inserted between the ApaI and KpnI (mTagBFP2) or PstI and KpnI sites (sYFP2). The pUC18R6K-mini-Tn7T-Gm plasmid [31] was a gift from Herbert Schweizer (Addgene plasmid # 65022; http://n2t.net/addgene:65022; RRID:Addgene_65022) The pFLP3 plasmid (Addgene #64946, [32]) was then introduced by conjugation to remove the FRT-flanked gentamicin resistance marker by FLP recombination. All strain constructions were confirmed with Sanger sequencing of the genomic region modified. All oligonucleotide primers are available in S2 Table.

### Fitness assays

Two types of competition fitness assays were performed following previously established protocols [8,11,14] except that flow cytometry was used to estimate the frequency of BFP vs. YFP cells in a population. We used a BioRad ZE5 Cell Analyzer flow cytometer to count 100,000 cells for each fitness assay replicate time point apart from *Psav* time 0 points, where we counted 200,000 cells, and gated on SSC-H and SSC-W to focus on single cells. We utilized the small particle optical detector, which measures forward scatter with a different laser that can resolve particles as small as 0.3 µm in diameter. A 488 nm laser combined with 530/30 filter was used for counting sYFP2 positive cells and the 405 nm laser combined with a 450/50 filter was used to quantify mTagBFP2 cells.

The first fitness assay measures the ability of a WS mutant to increase from low frequency in a dominant ancestral population. We intend this setup to be similar to the earliest stage of air-liquid interface colonization where a rare WS mutant attaches and grows at the surface with limited competition from other mutants. This is referred to here as “invasion fitness” and the assay was conducted by mixing the WS mutant with the wild type at a 1:99 starting ratio using a 1000-fold dilution of the wild type from an overnight culture. The assay was conducted under the same conditions as for the experimental evolution for 48 h.

In the second assay, we instead focus on “competition fitness” where a WS mutant competes 1:1 with a WS reference strain over 24 hours under the same conditions as for the experimental evolution starting with a 1000-fold dilution of overnight cultures. The reference strains have a WspA T293-T299 mutation in both *Psyr* and *Psav* and was chosen because it awas the most common mutation for both *Psyr* and *Psav*. We calculated selection coefficients (s) as previously described [33] s = [ln(R(t)/R(0))]/[t] where R is the ratio of the mutant to the reference and t is the number of generations of the entire population during the experiment. For both assays a value of s = 0 represents equal fitness, positive values increased fitness and negative values means decreased fitness relative to the reference strain. We quantified the fitness cost of the sYFP2 marker compared to mTagBFP2 marker in both the ancestral strains and the WspA reference strains and adjusted the selection coefficient based on this (*Psav* WT = +0.017, *Psav* WspA = +0.019, *Psyr* WT = −0.027, *Psyr* WspA = +0.024). Source data for all Fig 3 and Fig 4 are available in S3 Table.

### Motility assays

Motility assays were conducted to measure the swimming ability of the mutants compared to the ancestor. The assays were performed in TSBGM plates containing 0.3% agar (BD) and the diameter of the strains was measured after 24 hours of growth at room temperature. Each strain was tested in duplicates on two separate plates, resulting in four replicates.

### Enzymatic assay

Three hydrolytic enzymes, cellulase (Tokyo Chemical Industries Co. Ltd., Tokyo (Japan)), proteinase K (Thermo Scientific) and DNAse I (Thermo Scientific) were used to probe the structural basis of biofilm formation at the air-liquid interface. Clones derived from −80°C were streaked onto LB agar plates to obtain a single colony which was inoculated into 2 ml TSBGM and incubated at 25°C shaking overnight. Overnight cultures were diluted 1000-fold in 2 ml TSBGM with either cellulase (∼425 U (25 mg)), proteinase K (∼12 U (400 µg)), DNAse I (∼10 U (10 µl)) or no hydrolytic enzyme and incubated statically at 25°C. Biofilm formation was photographed at 24 h, 48 h and 72 h.

### Tn5 transposon mutagenesis

We used transposon mutagenesis of WS mutants to identify candidate genes required for the WS colony morphology [34] including putative structural components required for colonization of the air-liquid interface. The pCM639 plasmid containing the IS-Ω-kan/hah transposon was transferred by conjugation from *E. coli* SM10 *λpir* into WS mutants using helper plasmid pRK2013. We selected transconjugants on TSBGM plates supplemented with Congo Red, kanamycin and nitrofurantoin, for counter-selection of *E. coli*. Fewer than 1000 colonies from each independent conjugation were screened for loss of the WS colony morphology and reversion to an ancestral one. The transposons’ chromosomal insertions points were then determined by arbitrarily-primed PCR and Sanger sequencing [35].

### DNA sequencing

We sequenced the genomes of eleven WS mutants for *Psav* and identified mutations in *wspA* (PSPPH_3881), *wspE* (PSPPH_3877), *wspR* (PSPPH_3875), *wspC* (PSPPH_3879), PSPPH_0905 and PSPPH_2590. We also sequenced the genomes of four *Psyr* WS mutants that had mutations in *wspA* (PSPTO_1493), *wspE* (PSPTO_1497), PSPTO_3796 and *mwsR* (PSPTO_4631). Sanger sequencing of these candidate genes was then used to find the mutations causing the WS phenotype in the other mutants (S1 Table).

Genomic DNA was isolated with Genomic DNA Purification Kit (Thermo Fisher) for genome sequencing. For *Psyr*, sequencing libraries were prepared from 1μg DNA using the TruSeq PCRfree DNA sample preparation kit (cat# FC-121-3001/3002, Illumina Inc.). The library preparation was performed according to the manufacturers’ instructions (guide#15036187).

Sequencing was performed with MiSeq (Illumina Inc.) paired-end 300bp read length and v3 sequencing chemistry. Sequencing was performed by the SNP&SEQ Technology Platform in Uppsala. The facility is part of the National Genomics Infrastructure (NGI) Sweden and Science for Life Laboratory. The SNP&SEQ Platform is also supported by the Swedish Research Council and the Knut and Alice Wallenberg Foundation. For *Psav* we used a standard miniaturized protocol to prepare DNA libraries for sequencing using the NEBNext Ultra II FS DNA Library Prep Kit for Illumina (New England BioLabs Inc. (Ipswich, MA, USA)). The DNA libraries were quantified using a Qubit dsDNA HS kit and a Qubit 2.0 Fluorometer (Thermo Fisher Scientific Inc. (Waltham, MA,USA)). We used a MiSeq system (Illumina Inc. San Diego, CA, USA) to perform 250-bp paired end sequencing on the prepared libraries (Illumina MiSeq Kit V2 500 cycles).

Sequencing data were analyzed with using Geneious Prime software for Mac (v2022.0.2) with reads assembled against the *Psyr* (GenBank: genome AE016853.1, plasmids AE016854.1 and AE016855.1 [29] or *Psav* genome sequences (Genbank genome CP000058.1, plasmids CP000059.1 and CP000060.1, [30]). Complete sequencing data for these clones is available under NCBI BioProject PRJNA1075102. Sanger sequencing was performed by Eurofins Genomics and used to sequence candidate genes to find adaptive mutations and to confirm reconstructed mutations. Primer sequences are available in S2 Table.

## Author Contributions

**Conceptualization:** Peter A. Lind, Jennifer T. Pentz. **Formal analysis:** Jennifer T. Pentz, Peter A. Lind. **Funding acquisition:** Peter A. Lind. **Investigation:** Jennifer T. Pentz, Peter A. Lind, Aparna Biswas, Bassel Alsaed. **Methodology:** Jennifer T. Pentz, Peter A. Lind. **Resources:** Peter A. Lind. **Supervision:** Peter A. Lind. **Visualization:** Jennifer T. Pentz, Peter A. Lind. **Writing – original draft:** Jennifer T. Pentz, Peter A. Lind. **Writing – review & editing:** Jennifer T. Pentz, Aparna Biswas, Peter A. Lind.

## Acknowledgements

This work was supported by the Kempe foundations (SMK-1858.1), Carl Trygger’s Foundation for Scientific Research (CTS 19:204), O.E och Edla Johanssons vetenskapliga stiftelse, Åke Wiberg’s foundation (M18-0142) and Magnus Bergvall’s Foundation [P.A.L]. The funders had no role in study design, data collection and analysis, decision to publish, or preparation of the manuscript. This work has been cleared for public release by the Los Alamos National Laboratory LA-UR-24-21205.

---

**S1 Fig.**
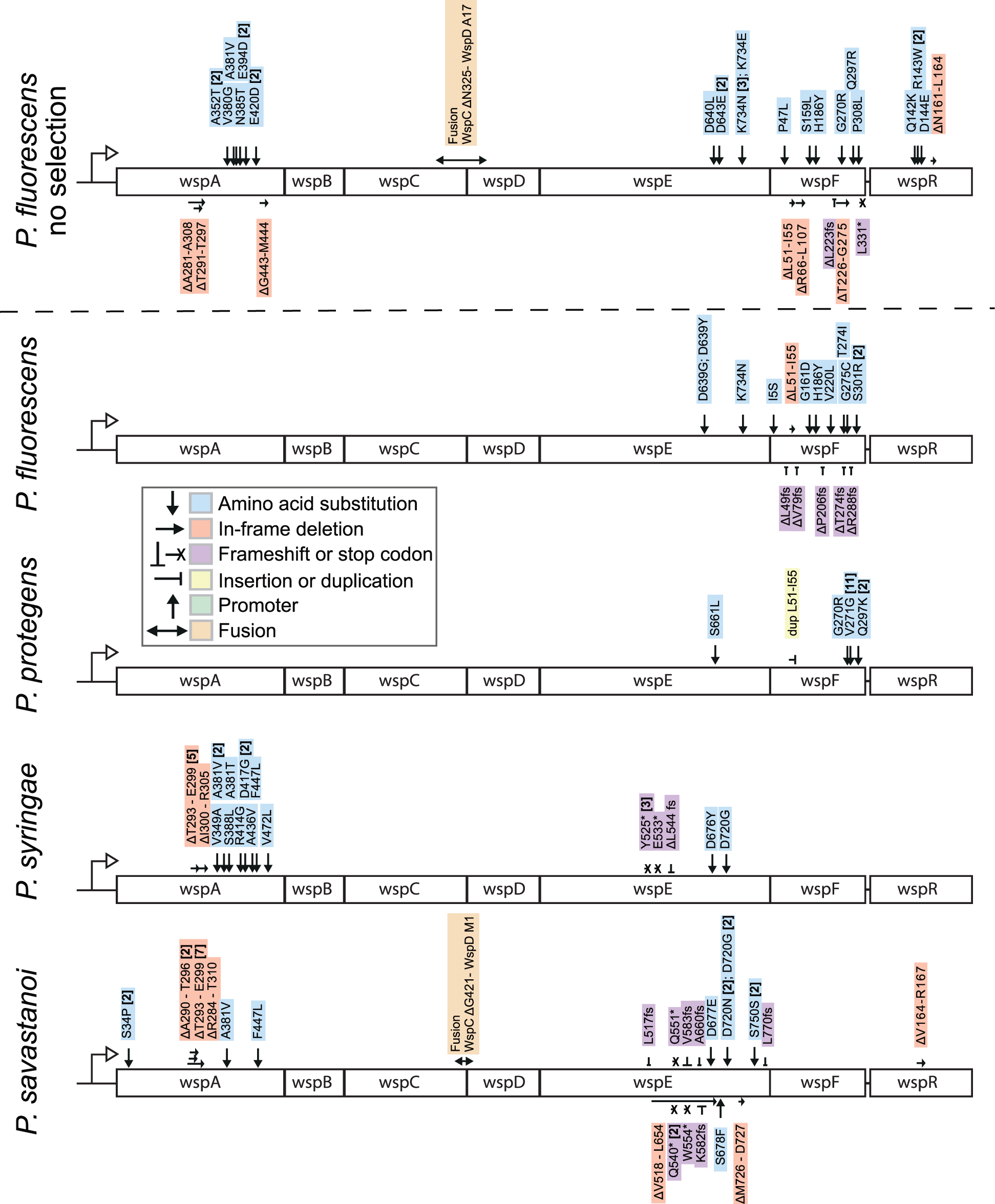
Comparison of WS mutations in the *wsp* operon. Data for “*P. fluorescen*s no selection” is from Lind *et al.* 2019 [7] where a reporter construct was used to isolate WS mutants from a fluctuation assay without selecting for ability to colonize the air-liquid interface. The other mutants were isolated after experimental evolution with data from McDonald *et al.* 2009 [10] for *P. fluorescens*, Pentz *et al.* 2021 [8] for *P. protegens* and this work for *P. syringae* and *P. savastanoi*.

**S2 Fig.**
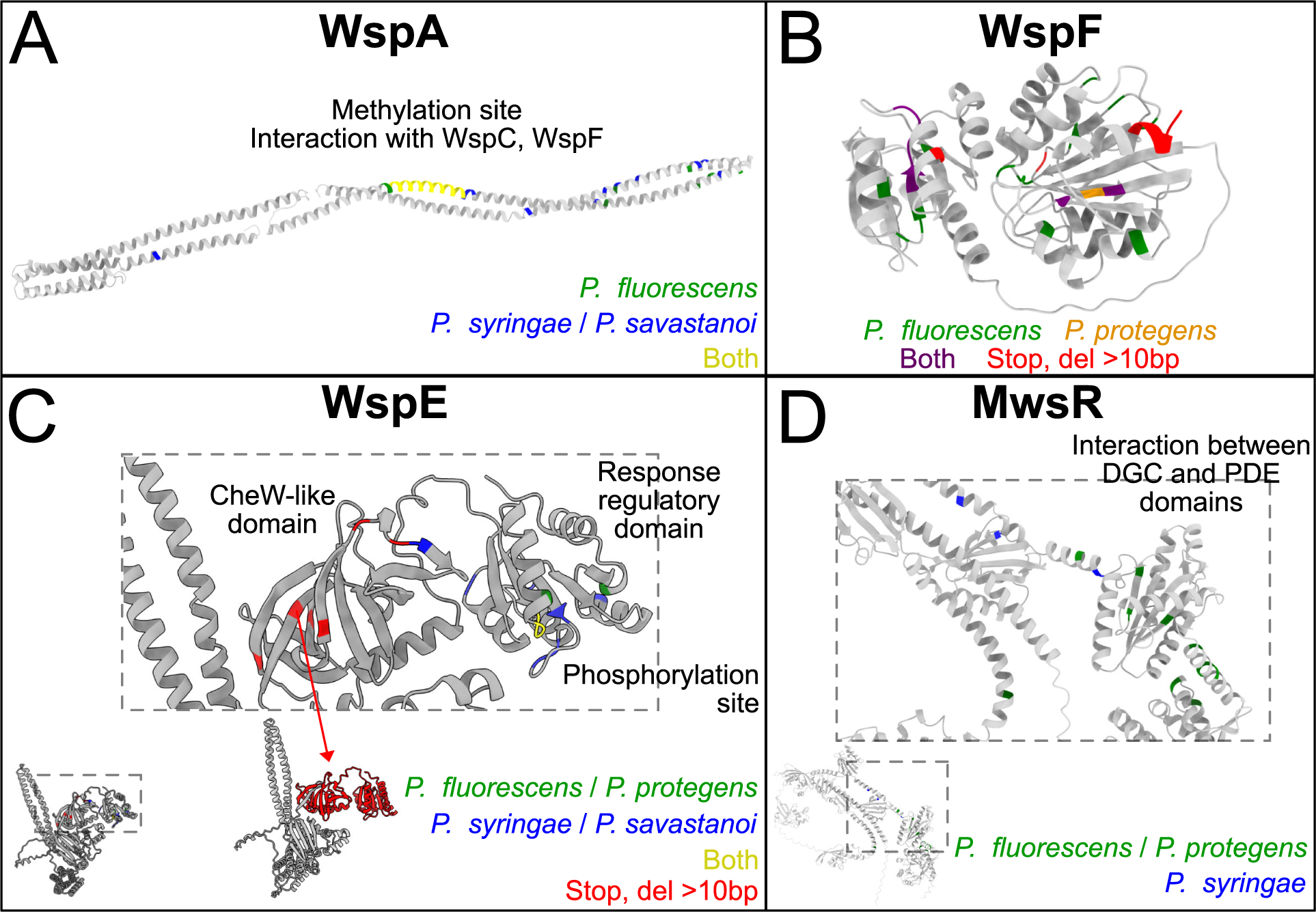
Comparison of WS mutations in homology models of protein structures for WspA, WspF, WspE and MwsR. All mutations shown in Fig. S1 from this work, Lind et al 2019 (Pflu no selection), McDonald et al 2009, and Pentz et al 2021 mapped onto homology models for WspA, WspF, WspE, and MwsR. (A) WspA mutants are similar for both *Pflu* and *Psyr*/*Psav* and occur at the methylation site as previously predicted (Pentz et al 2021). (B) WspF mutants were only found in *Pflu*/*Ppro* and are distributed across the whole protein as any inactivating mutation will result in WS. (C) WspE mutants were found in both *Pflu*/*Ppro* and *Psyr*/*Psav*. *Pflu*/*Ppro* had specific point mutations at an active site, while *Psyr*/*Psav* had mutations that delete the whole CheW-like domain (inset shows deleted region of WspE in red for mutation Y525*). (D) MwsR mutations mostly occurred at the interaction between the DGC and PDE domains, as predicted in Pentz et al 2021.

**S3 Fig.**
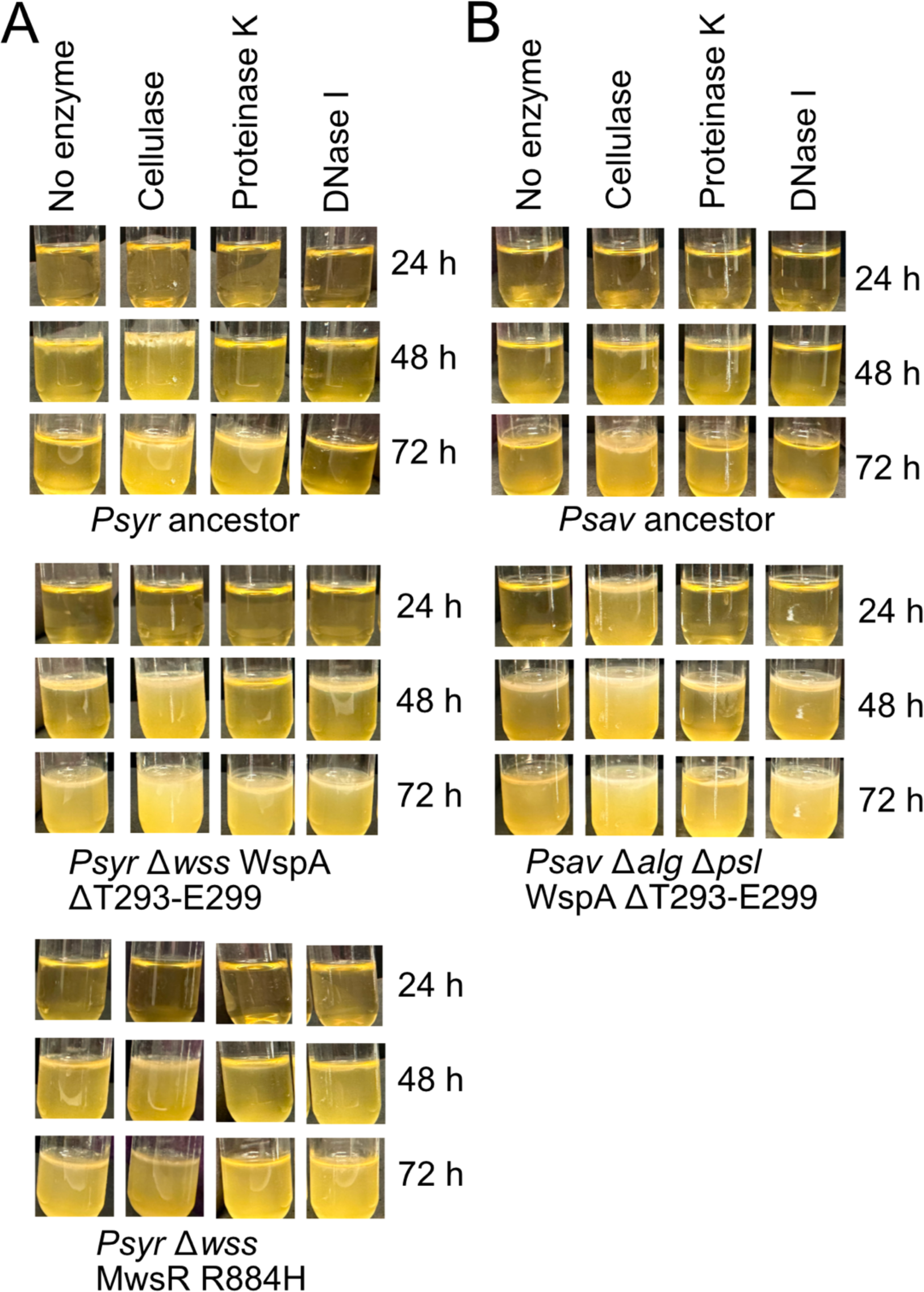
Biofilm formation under static growth with cellulase, proteinase K and DNase I. (*A*) *Psyr* ancestor and Psyr WS mutants with deletion of the *wss* operon, encoding the genes for cellulose biosynthesis. (*B*) Psav ancestor and WspA mutant with deletion of *alg* and *psl* operons.

